# The 3’ Region of the ZPA Regulatory Sequence (ZRS) is required for activity and contains a critical E-box

**DOI:** 10.1101/2025.02.01.636055

**Authors:** Kathryn F. Ball, Stephen Manu, Abbie K. Underhill, Sarah R. Rudd, Madison M. Malone, Japhet Amoah, Allen M. Cooper, Charmaine U. Pira, Kerby C. Oberg

## Abstract

1

**Background:** During development, Hand2 and Hoxd13 transcription factors (TFs) regulate Sonic hedgehog (Shh) expression in the zone of polarizing activity (ZPA) in the distal posterior limb mesoderm. The ZPA regulatory sequence (ZRS) is a conserved, limb-specific enhancer that controls Shh expression. The ZRS can be divided into 5’, central, and 3’ subdomains, each with an E-box site that can bind basic helix-loop-helix (bHLH) TFs like Hand2. In addition, two Hoxd13 sites are present in the 5’ and central subdomains. Hand2 purportedly binds the ZRS through the central E-box, and both Hand2 and Hoxd13 have been shown to activate the ZRS *in vitro*. We hypothesized that the central E-box was required for activity, while the other E-boxes and Hoxd13 sites localize ZRS activity to the distal posterior limb mesoderm.

**Methods:** To identify the functional role of each subdomain, we generated three ZRS fragments (5’, central, and 3’) and combined fragment constructs to test subdomain collective contributions. Additionally, we disrupted the five binding sites, alone or in concert, using site-directed mutagenesis. All ZRS constructs were cloned into a GFP reporter and evaluated in an *in vivo* chicken limb bioassay. We validated our findings using select ZRS constructs in transgenic mice.

**Results:** We found that the 3’ fragment was necessary for ZRS activity, while the 5’ and central fragments had no activity alone or when combined. However, combining the 3’ fragment with the 5’ fragment restored robust activity. Further, mutation of all five binding sites markedly reduced ZRS activity. Reinstating each of the Hoxd13 sites restored focal activity, while restoring the 5’ and central E-boxes had little effect. However, the 3’ E-box proved sufficient for robust activity even in the absence of the other four binding sites.

**Conclusions:** Our data indicate that the ZRS 3’, not the central, subdomain is necessary for activity and contains the 3’ E-box that Hand2 likely uses to induce Shh expression, while the 5’ and central E-boxes appear to be inhibitory. Our data also suggest that the Hoxd13 binding sites promote localized activity within the ZPA.

## 2 Introduction

Sonic hedgehog (Shh) is a secreted signaling factor that directs morphogenesis in several organs during development including the neural tube, early gut, and limb. The zone of polarizing activity (ZPA) refers to a small subpopulation of mesenchymal cells in the posterior distal aspect of the developing limb that secrete Shh to direct anterior-posterior (AP) patterning. Shh knock-out (KO) in mice results in loss of posterior limb structures such as the ulna and fibula in the zeugopod, and all but a single digit in the autopod (Chiang et al., 2001; Kraus et al., 2001). Despite Shh’s pivotal role in limb patterning, the mechanisms that maintain its expression within the ZPA during progressive limb outgrowth remain unclear.

*Cis*-regulatory modules (CRMs) are DNA sequences that sense cellular cues for tissue-specific transcription factors (TFs). The ZPA regulatory sequence (ZRS) is a limb-specific CRM located approximately one million bases upstream of the *Shh* promoter within intron 5 of the *Lmbr1* gene. The ZRS is necessary for Shh expression in the ZPA, as demonstrated by the loss of Shh expression after a spontaneous ZRS microdeletion in chickens (Ros et al., 2003) or after ZRS KO in mice (Sagai et al., 2005). The ZRS can be divided into three subdomains that are conserved across vertebrate species, hereafter called: 5’, central, and 3’ (Figure 1A, purple boxes). Investigating ZRS architecture can help identify themes in CRM function and elucidate the overarching principles of regulatory DNA.

**Figure 1.**
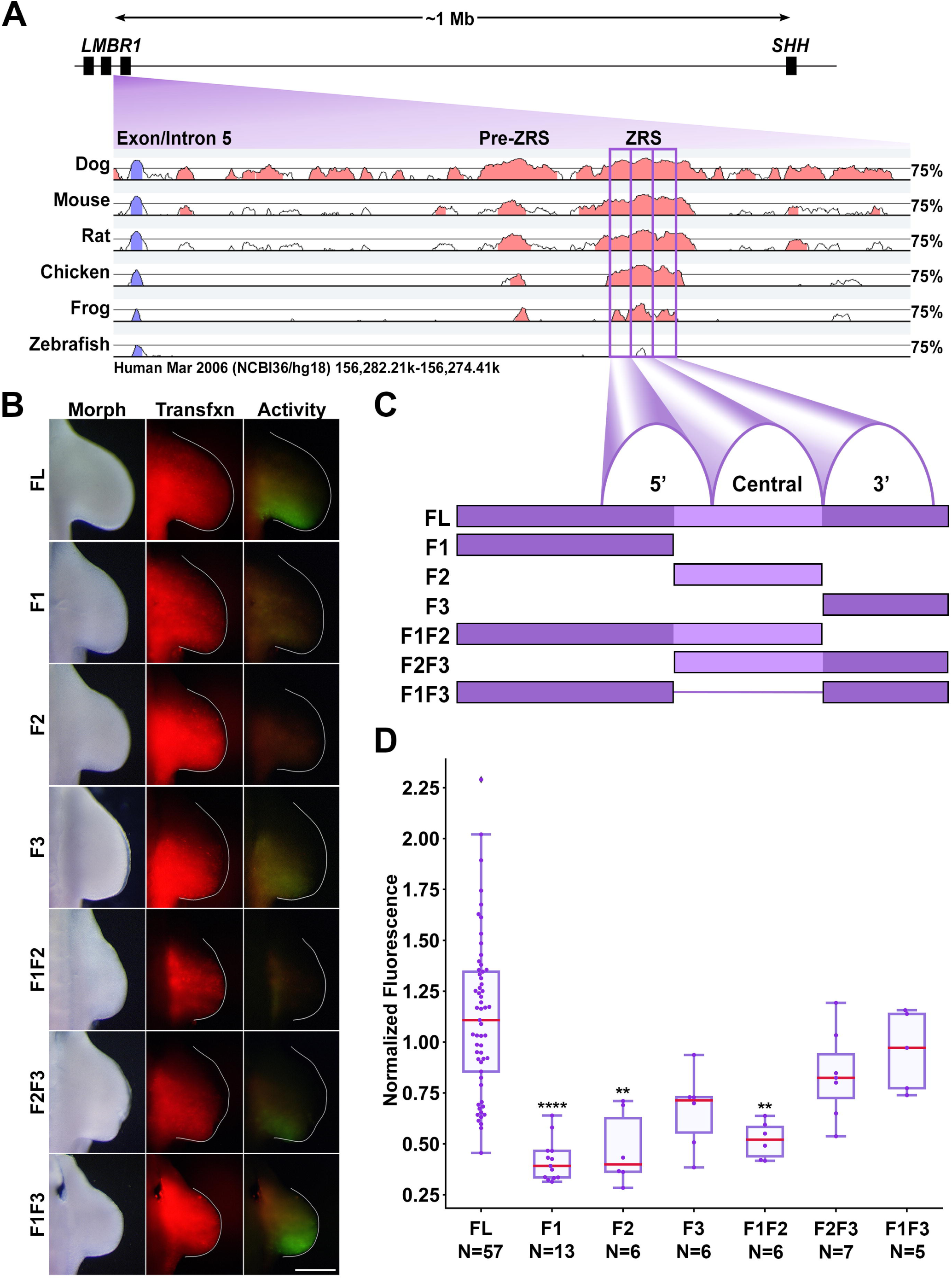
The 3’ subdomain of the ZRS is required for activity. A) Diagram of the ZRS locus relative to Shh and pairwise conservation of each listed species in comparison to the human sequence (VISTA point). The ZRS subdomains are boxed in purple. B) Activity of ZRS and subdomain fragments. Morph: morphology, Tfxn: transfection control. Images are dorsal view with top: anterior, right: distal. Scale Bar = 1mM. C) Diagram of the conserved ZRS fragments in this study. FL: full length, F1: Fragment 1, F2: Fragment 2 etc. D) Box- and swarm plots of fragment activity (GFP intensity) normalized to transfection control (RFP intensity). Changes in activity were compared with a Kruskal-Wallis test followed by Dunn’s test. ** = p < 0.01, **** = p < 0.0001, ns = not significant. N refers to the number of embryos per group. Experimental groups were repeated in at least three independent experiments.

The transcription factors Hoxd13 and Hand2 regulate Shh expression in the limb. Hoxd13 contributes to AP polarity, and early anterior Hoxd13 misexpression results in anterior Shh expression (Zakany et al., 2004). Hand2 is necessary for Shh expression; Hand2-deficient mouse limb buds display a phenotype similar to Shh loss-of-function limbs (Galli et al., 2010). Conversely, anterior Hand2 misexpression in the limb bud produces ectopic Shh expression leading to mirror-image digit duplication (Charite et al., 2000). Hoxd13 and Hand2 bind the ZRS and each other; they can also independently transactivate ZRS-luciferase *in vitro* and, when combined, can transactivate ZRS synergistically (Capellini et al., 2006; Galli et al., 2010).

Hand2 is a basic helix-loop-helix (bHLH) TF that forms homo- and heterodimers with other bHLH factors such as Hand1, Twist1, E12, and E47 (Dai and Cserjesi, 2002; Firulli et al., 2005; Koyano-Nakagawa et al., 2022). Two bHLH monomers must dimerize to form a functional transcription factor, and since each monomer contributes its DNA binding domain to make half of the whole DNA binding region, dimer composition can affect the affinity to a binding site (Dai et al., 2002; Firulli et al., 2007). Even small changes in binding affinity can result in a pathological phenotype (Lim et al., 2024).

Transcription factors in the bHLH family bind E-boxes, hexamers with a core “CANNTG” motif. An E-box with Hand2’s consensus binding sequence (CAGATG) in the central ZRS subdomain is purported to be the Hand2 binding site (Galli et al., 2010; Osterwalder et al., 2014). Other factors including Snail and Slug, zinc-finger TFs, and Hey1 are expressed in the early limb and could also bind this E-box. We set out to interrogate this E-box along with two others within the highly conserved ZRS to determine their relevance to ZRS activity.

Efforts have been made to map the ZRS; however, this work is incomplete. Characterizing the ZRS transcription factor binding site (TFBS) landscape is critical to both understanding development and clinical Shh dysregulation. More than 30 single-nucleotide variations (SNVs) within the ZRS have been documented, most of which result in preaxial polydactyly (PPD) and/or triphalangeal thumb (TPT) (Supplementary Table 1). A majority of the human SNVs (22/34) are located within the central ZRS subdomain suggesting this region is susceptible to perturbation (Supplementary Figure S1). In this study, we demonstrate the 3’ E-box is critical for ZRS activation, the 5’ and central E-boxes are repressive, and the Hox sites are activating.

## 3 Materials and Methods

### Plasmid Construction

To test ZRS activity *in ovo*, we generated expression constructs with ptk-EGFP plasmid (a gift from Dr. Masanori Uchikawa, Osaka University, Japan) (Uchikawa et al., 2003), which contains the minimal HSV TK promoter linked to an enhanced GFP reporter gene. Chicken ZRS (cZRS, a 1,373 bp fragment, Assembly IDs UCSC: GRCg6a/galGal6 and NCBI: 1668981, Chr2:8,553,160-8,554,532) or human ZRS (hZRS, a 1,198bp fragment, Assembly IDS UCSC Hg38 and NCBI: GRCh38.p14, Chr7:156,791,072-156,792,269) was isolated by PCR from genomic DNA and ligated into pTK-EGFP at the XhoI restriction site. Constructs containing individual conserved peaks were generated through progressive 3’ digestion with the Erase-a-Base system (Promega, Madison, WI). pCAGGS-RFP plasmid (a gift from Dr. Cheryl Tickle, University of Dundee, Scotland) (Das et al., 2006) was co-electroporated to verify transfection. Plasmids were isolated and purified using the EndoFree Plasmid Maxiprep Kit (Qiagen, Valencia, CA).

### Site-Directed Mutagenesis

To disrupt transcription factor binding, we altered three-to-four core bases of each putative binding site with the QuikChange Multi Site-Directed Mutagenesis Kit (Agilent Technologies, Santa Clara, CA) while also introducing a restriction site for screening. Mutant sequences were analyzed with CiiiDER (Gearing et al., 2019) to ensure no new binding sites relevant to limb development were introduced. NEB5-α competent cells were transformed with mutated constructs. Transformants were screened using the new restriction site and constructs were confirmed via Sanger sequencing (Eton Bio, San Diego, CA). All genomic and mutagenic primers are listed in Supplementary Table 2.

### Targeted Regional Electroporation (TREP)

Chicken embryos were staged according to the Hamburger and Hamilton (HH) method (Hamburger and Hamilton, 1951). The embryonic coelom of the lateral plate mesoderm of stage HH14 embryos was injected with DNA solution (2 µg/µL pTK-ZRS-EGFP, 0.2 µg/µL pCAGGS-RFP with Fastgreen and Tris-EDTA buffer). Plasmids were electroporated into the presumptive limb using the CUY-21 Electroporator (Protech International Inc., Boerne, TX) as previously described (PIRA et al., 2008). Embryos were incubated for 48 hours post-electroporation then harvested. We visualized fluorescence with a Leica MZ FLIII fluorescence stereo microscope using 41012 HQ:FLP FITC/EGFP and 10446365 TXR filters (Chroma Technology Corp., Brattleboro, VT); images were captured with a Sony DKC-5000 camera and acquired using *Adobe Photoshop* (version 6.0).

### Image Analysis

Image analysis was performed using a workflow written in *Python* (3.9.12). In short, images were converted to grayscale, passed through a bilateral denoise filter, then the region of ‘Limb’ was determined using a combination of Otsu thresholding and manual input (to separate limb from body wall) on the light image. To limit RFP measurement to relevant tissue only, three different masks were made for each limb: the Limb mask excluded background and non-limb tissue, the Posterior mask excluded tissue that might be transfected, but would not express wild-type activity, and the ZPA mask that limits measurement to the region of active *Shh* transcription. Diagrams of the masks can be seen in Figure 2C. We used the Posterior mask for all image analyses in this paper except for the Hoxd13 mutant analysis shown in Figure 3C. The region of transfection was determined using the masked RFP image and Otsu thresholding. The region of enhancer activity was determined by applying the ‘RFP’ mask to the GFP image combined with Otsu thresholding. Pixel number and intensity were measured within the appropriate mask (RFP on the RFP image, GFP on the GFP image), and relative enhancer activity was determined by normalizing total GFP intensity to total RFP intensity. This normalization accounts for differences in transfection. A *Jupyter* notebook of the code used is available at https://github.com/KateBall/Quantitative_Image_Analysis under the GNU Public License (GPL, ver. 3). A preprint describing the method in detail can be found at **[BioRxiv Reference]**.

**Figure 2.**
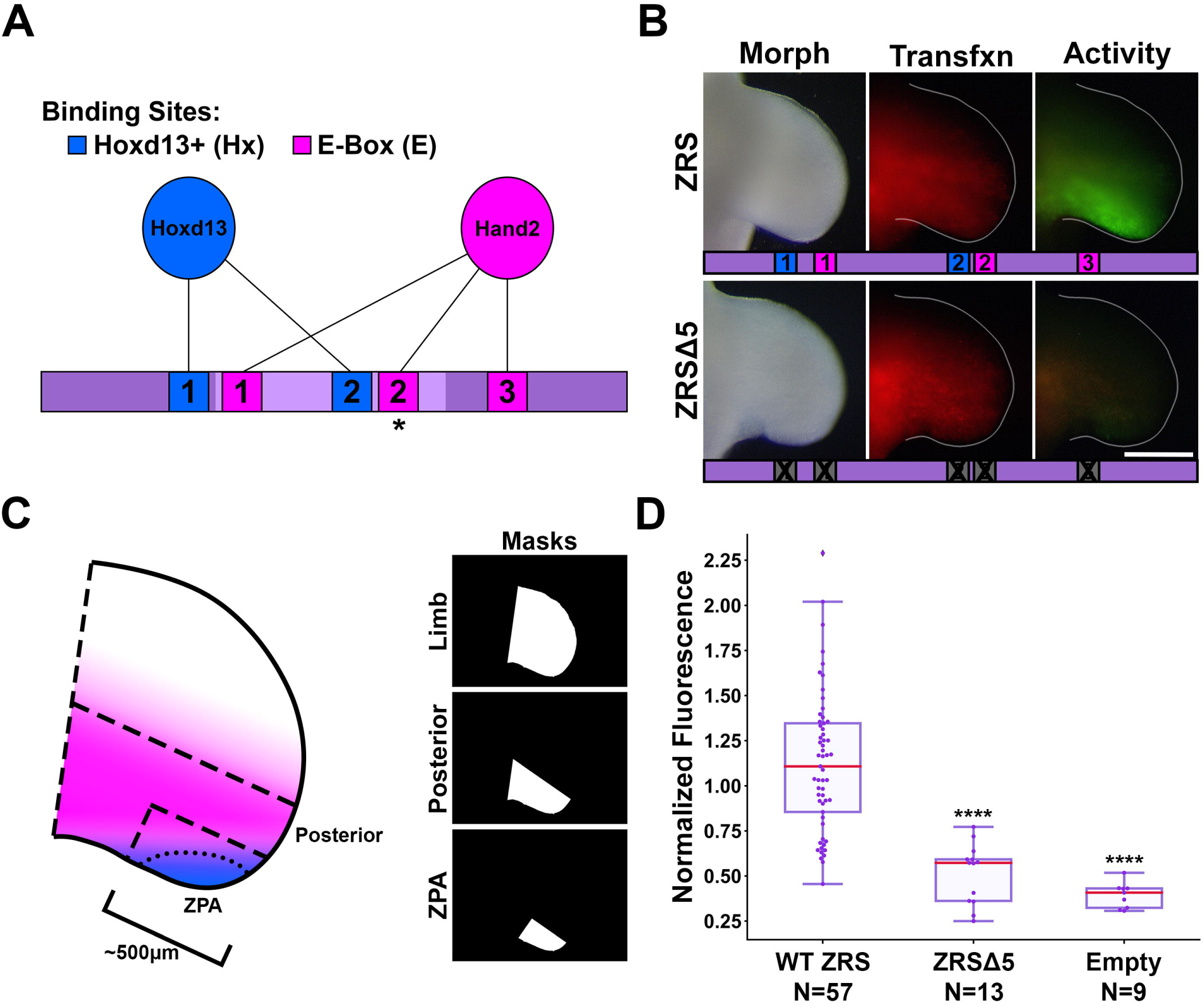
Loss of Hoxd13 binding sites and E-boxes reduces ZRS activity. A) Diagram of key transcription factors and binding sites in this study. Asterisk indicates reported Hand2 binding site. B) Activity of wild type ZRS (WT) or with five binding sites mutated (ZRSΔ5). C) Diagram showing Hand2 (pink) and Hoxd13 (blue) expression pattern overlap. Masks indicate the three regions in which fluorescence was measured and correspond to dashed regions on the limb diagram. D) Box- and swarm plots of reporter activity (GFP intensity) normalized to transfection control (RFP intensity). Data collected using the Posterior mask. Changes in activity were compared with a Kruskal-Wallis test followed by Dunn’s test. **** = p < 0.0001, ns = not significant. N refers to the number of embryos per group. Experimental groups were repeated in at least three independent experiments.

**Figure 3.**
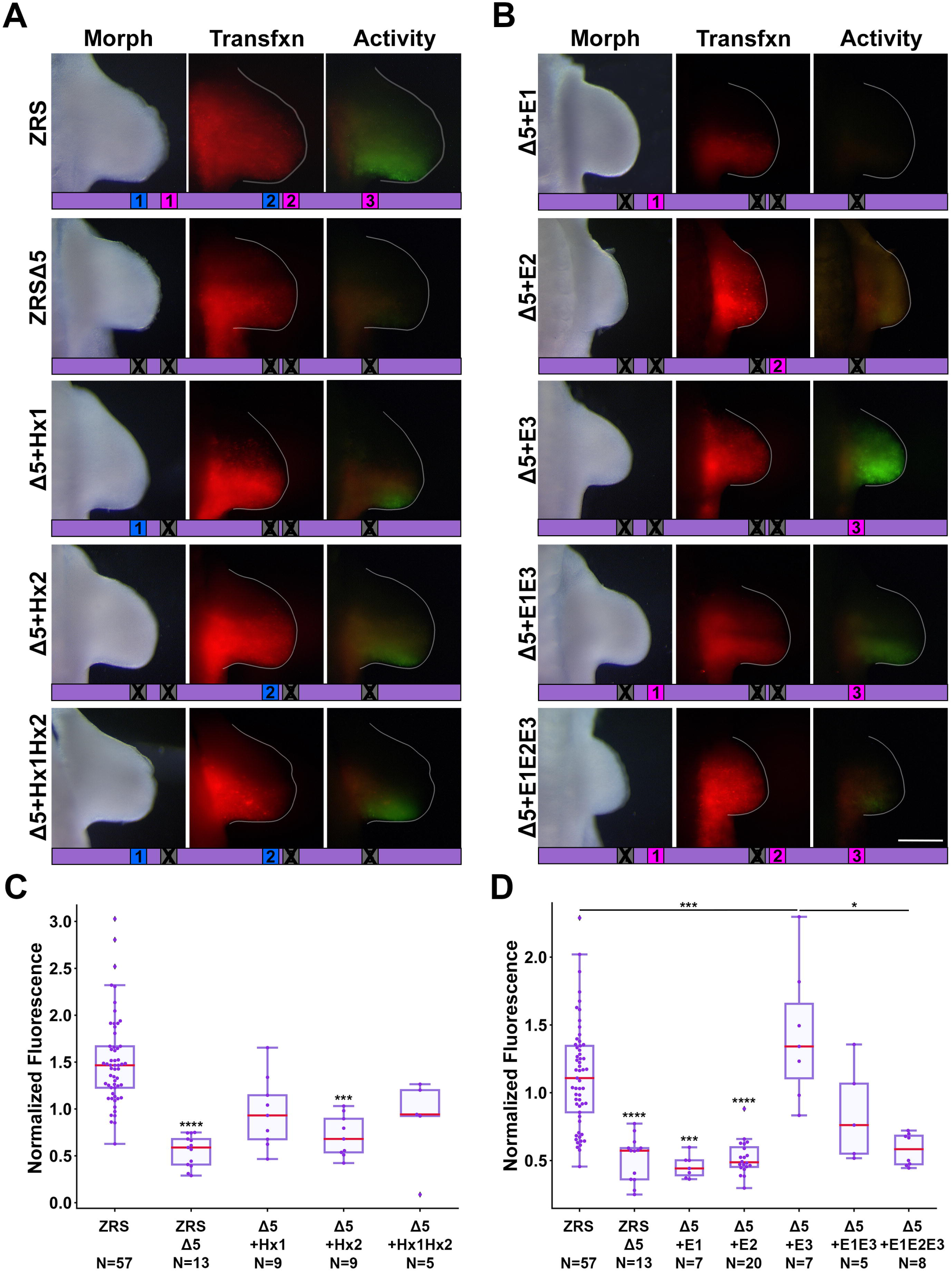
E-box and Hoxd13 binding sites differentially regulate ZRS activity. A) Activity of ZRS: wild type (WT) or with five binding sites mutated (ZRSΔ5), restoration of each Hoxd13 site, alone and in concert. B) Restoration of each E-box, alone and in concert. Diagrams below each image limb show which binding sites are present or absent in the given construct. C, D) Box- and swarm plots of reporter activity (GFP intensity) normalized to transfection control (RFP intensity). Activity was measured using the ZPA mask in C) and the Posterior mask in D). Changes in activity were compared with a Kruskal-Wallis test followed by Dunn’s test. * = p < 0.05, *** = p < 0.001, **** = p < 0.0001, ns = not significant. N refers to the number of embryos per group. Experimental groups were repeated in at least three independent experiments.

### Transgenic Mice

Human ZRS (hZRS) and its Δ5 mutant (hZRSΔ5) were cloned into the HSP68-LacZ plasmid kindly provided by Dr. Nadav Ahituv, UC San Francisco, CA (Pennacchio et al., 2006). The constructs were used to generate transgenic mouse embryos (Cyagen transgenic service, Santa Clara, CA). Embryos were harvested at e12.5 and processed for detection of LacZ activity.

### Single-cell RNA Sequencing Analysis

Mouse forelimb single-cell RNA sequencing data from (He et al., 2020) was analyzed using Partek® Flow® software, v10.0.23.0720. We imported the h5 matrices and filtered out cells with less than 600 transcripts or with 10% or more reads mapping to the mitochondrial genome. We normalized gene expression using the standard equation: E_x,y_ = log2[(CPM_x,y_) + 1], where CPM_x,y_ refers to counts per-million for gene x, in sample y. Genes that were not detected in more than three cells were also filtered out. Differential expression analysis was performed on Shh-expressing (Shh+) cells (normalized expression greater than 0.5) versus Shh non-expressing (Shh-) cells (normalized expression lower than 0.5) using Kruskal-Wallis. p-values were adjusted using the Bonferroni method.

### Binding Site Affinity Analysis

Relative binding affinity was calculated using protein binding microarray data from the UniProbe database (http://thebrain.bwh.harvard.edu/uniprobe/index.php) (Hume et al., 2015) and processed using *Python* code adapted from (Lim et al., 2024). The adapted code is available at (https://github.com/KateBall/ZRS-2025). For each transcription factor, the relative affinity values represent a ratio of the median intensities of the given 8mer over the factor’s optimal 8mer from the PBM data.

### Statistical Analysis

TREP data were collected over at least three separate experiments per group. The interquartile ranges are shown via box plots with medians in red; individual data points, each corresponding to the limb of a separate embryo, are shown via swarm plot. The reported sample sizes (N) on each plot correspond to the number of embryos in each group. Note that the same data for chicken wild-type ZRS (referred to as “FL” and “WT”) and Empty vector (“Empty”) are shown in both Figures 1 and 2. Quantitative data were analyzed with the *Python* modules *Pandas*, *NumPy*, *SciPy*, and *scikit_posthocs*, and were visualized with *Matplotlib* and *Seaborn*. To check for normality, we visually inspected the data using histograms and used the Shapiro-Wilk test for normality. Outliers were identified using the interquartile range method, but not dropped as it would necessitate removing some of the smaller groups from analysis. Statistically significant (p<0.05) differences were determined with the Kruskal-Wallis test followed by Dunn’s multiple comparisons test with Bonferroni p-value correction or the Mann-Whitney U test when only two groups were compared. * = p< 0.05; ** = p< 0.005; *** = p< 0.0005; **** = p< 0.0001. A *Jupyter* notebook of the complete statistical analysis can be found at (https://github.com/KateBall/ZRS-2025).

#### Note

All figures were made using *Adobe Photoshop (CC)*. Subfigures containing plots, masks, or limbs with contours also used *Matplotlib 3.5.1*, *Seaborn 0.12.2*, and *Statannotations 0.4.4*. Original psd files are available upon request. Any alterations made and are intended for clarity and aesthetic purposes only. All quantitative data used in this study are collected from raw, unaltered image files.

## 4 Results

### The ZRS 3’ region is required for activity

To uncover the regulatory role of the ZRS subdomains, we divided the 1373 bp chicken ZRS into 5’ (749 bp), central (309 bp), and 3’ (302 bp) fragments and generated reporter constructs containing single and combined fragments (Figure 1). The 3’ fragment (F3) demonstrated activity consistent with full-length (FL) wild type; while the 5’ and central fragments (F1 and F2, respectively) had significantly less activity compared to FL. The lack of activity in the central fragment was surprising because Hand2 reportedly binds the central E-box (E-box 2) (Osterwalder et al., 2014). This shifted our attention to the 3’ fragment since it retained activity consistent with a critical role in ZRS activation.

The combination of the 3’ fragment with the 5’ fragment (F1F3) exhibited more intense activity than the 3’ fragment alone (Mann-Whitney U Test p = 0.017) and was similar to wild type in both pattern and intensity. The construct combining the 5’ and central portions of the ZRS (F1F2) still lacked statistically significant activity when compared to the wild type ZRS. We also tested the F2F3 fragment, and did not find a significant increase over F3. These data indicate that the 5’ (F1) subdomain contributes to activity, but only in the presence of the 3’ region (F3).

### Loss of Hoxd13 binding sites 1 & 2 and E-boxes 1-3 nearly abolishes ZRS activity

Since Hand2 and Hoxd13 have been shown to work together to activate the ZRS *in vitro* (Galli et al., 2010), we set out to identify probable binding sites for each. While Hox TFs are known for their binding site promiscuity, Hoxd13 has a high affinity to C/TAATAAAA motifs, and we identified and interrogated two sites favorable for Hoxd13. We evaluated three E-boxes within the highly conserved ZRS region that are found in human, mouse, and chicken (Figure 2A). E-boxes 1, 2, and 3 are in the 5’, central, and 3’ subdomains, respectively. E-box 2 (CAGATG) in the central subdomain is Hand2’s predicted binding motif (for more detail see Supplementary Figure S2). We mutated all five binding sites in concert (ZRSΔ5) as a screening process to see if any of the sites had functional relevance and found that loss of all five binding sites nearly abolished ZRS activity (Figure 2).

### The presence of at least one functional Hoxd13 binding site in the ZRSΔ5 restores focal activity

To determine the relative contribution of each binding site to ZRS activity, we used the ZRSΔ5 as a baseline and restored each binding site individually and with the others of its class. With this assay, the signal measured is the result of accumulated GFP within cells having a history of ZRS activation over the 48-hour incubation period. WT activity is the result of early ZRS induction, presumably from Hand2, which is expressed prior to limb outgrowth, and maintained by Hoxd13 a few stages later in the limb bud. The presence of at least one Hoxd13 binding site (ZRSΔ5+Hx1, ZRSΔ5+Hx2, or ZRSΔ5+Hx1Hx2), produced activity that appears more focal than wild type (Figure 3). To capture the differential activity in the ZPA domain, we used a ZPA mask to quantitate fluorescence (Figure 2C). We found that within the ZPA domain, ZRSΔ5+Hx1 recovered activity with intensity that was not significantly different from wild type, though ZRSΔ5+Hx2 was less (Figure 3C). Thus, the focal activity of the ZRSΔ5+Hx1 and ZRSΔ5+Hx2 constructs may reflect a late maintenance-related activation with reduced GFP accumulation. Interestingly, F1F2 has no activity despite containing both Hoxd13 binding sites, suggesting that other binding sites in the 3’ region are necessary to support Hoxd13–related activation.

### E-box 3, and not the canonical Hand2 binding site, restores ZRS activity

Restoring E-box 1 or E-box 2 in ZRSΔ5 did not significantly increase activity (Figure 3B). However, restoring E-box 3 in the context of ZRSΔ5 produced activity significantly greater than wild type, suggesting the 3’ E-box drives ZRS activity and may be the site Hand2 uses to activate the ZRS. Surprisingly, ZRS activity in the presence of the 3’ E-box 3 in combination with E-boxes 1 and 2 (ZRSΔ5+E1E2E3) results in a reduction of activity (Figure 3D), indicating E-boxes 1 and 2 perform an inhibitory role. The addition of E-box 1 (ZRSΔ5+E1E3) reduces activity compared to ZRSΔ5+E3, but not to a significant degree.

E-box 3 restores ZRS activity despite the absence of two Hoxd13 sites. This may be possible because other Hox binding sites are present in the ZRS and Hox TFs are known to be promiscuous. Thus, it is possible that Hoxd13 is acting on ZRS through other Hox binding sites. We initially suspected that loss of the purported Hand2 binding site, E-box 2, would be sufficient to eliminate ZRS activity. However, others have shown that the central ZRS subdomain (F2) is dispensable for activity (Lettice et al., 2017). We also found that the ZRS maintained activity following site-directed mutagenesis of the central E-box alone, consistent with Lettice and colleagues (Supplementary Figure S3).

### The conserved Hoxd13 and E-Box sites are also critical for human ZRS activity

To determine whether the necessity of the five binding sites is conserved across species, we repeated the *in vivo* bioassays using human ZRS (hZRS) and found that the absence of the five binding sites (hZRSΔ5) also depleted hZRS activity (Figure 4A-B). We then interrogated hZRS in the transgenic murine model using a β-galactosidase assay that results in blue precipitate in locations that have had ZRS activity. Although both wild type and hZRSΔ5 showed evidence of ZRS activity (Figure 4C), the wild type hZRS had blue precipitate encompassing digital rays 4 and 5. However, hZRSΔ5 activity was substantially reduced and restricted to the distal tips of digital rays 4 and 5, indicating the five binding sites are needed for normal ZRS activity. The hZRSΔ5 construct resulted in some activity (arrowhead) outside of the ZPA-related region stained by the wild type ZRS, despite verifying that no new binding sites were introduced. Interestingly, the wild-type construct also produced some anterior ectopic staining (Supplementary Figure S4). This may be due to the nature of random genomic insertion or be a function of the ZRS being out of context with other CRMs in its regulatory landscape that provide additional anterior inhibition (Petit et al., 2016).

**Figure 4.**
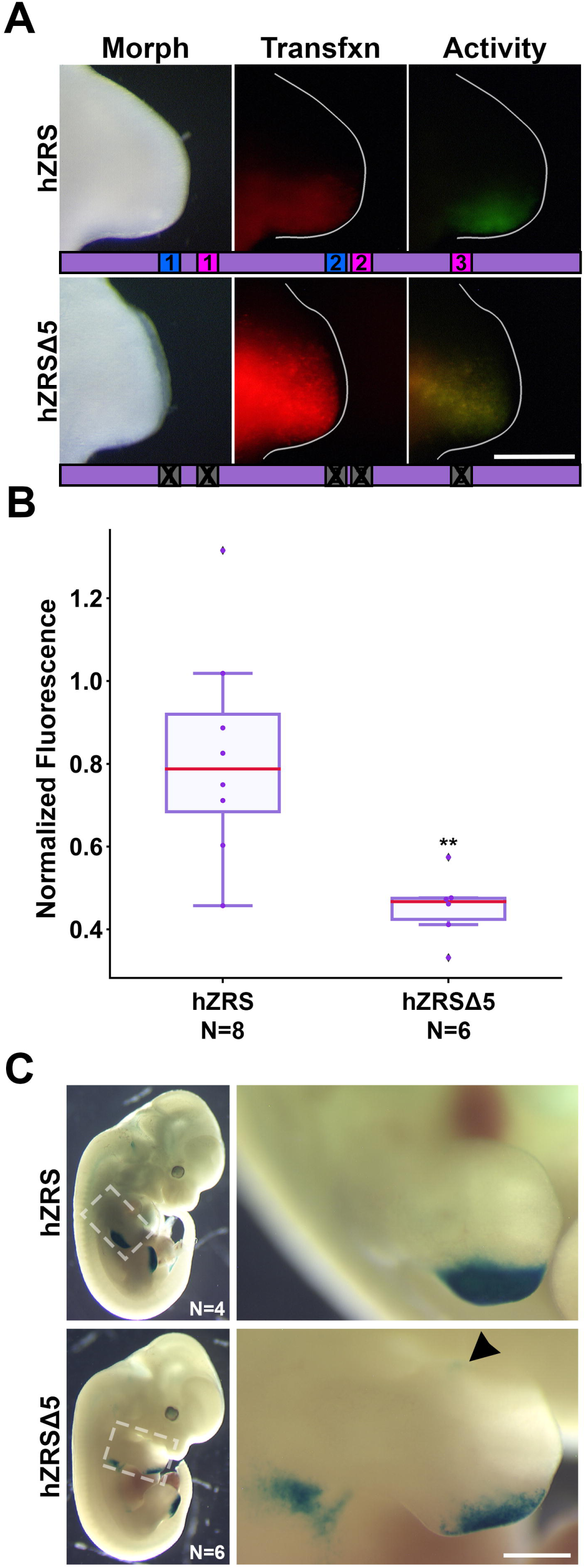
The importance of the five binding sites is conserved in human ZRS. A) Activity of wild type human ZRS (hZRS) or with five binding sites mutated (hZRSΔ5). B) Box- and swarm plots of reporter activity (GFP intensity) normalized to transfection control (RFP intensity). Changes in activity were compared with a Mann-Whitney U test. ** = p < 0.01. N refers to the number of embryos per group. Experimental groups were repeated in two-three independent experiments. C) hZRS and hZRSΔ5 activity in mouse embryos harvested at e12.5. Arrowhead points to ectopic anterior activity. Size bar represents 1mm.

### Identification of additional potential ZRS cofactors in Shh-expressing cells

To identify any additional potential cofactors involved in ZRS regulation, we analyzed published single-cell RNA sequencing data from mouse embryo limbs (He et al., 2020). We performed differential analysis on Shh+ versus Shh-cells from embryonic day (e)10.5 and e11 mouse limbs (Figure 5A and Figure 5B, respectively). Several Shh-related genes were differentially expressed in an expected pattern supporting the analysis (e.g., Hand2 and Hoxd13 upregulation at e10.5 and e11), as well as other hox factors (Hoxd12, Hoxd11, and Hoxa11). We identified several additional bHLH TFs (Hey1, Snail1, Twist2) upregulated at e10.5 or e11 that could function as co-factors with Hand2 in ZRS regulation (Supplementary Table 3). When we analyzed e12 limb bud cells, most differential expression had dropped off likely due to decreased Shh expression (Supplementary Figure S5).

**Figure 5.**
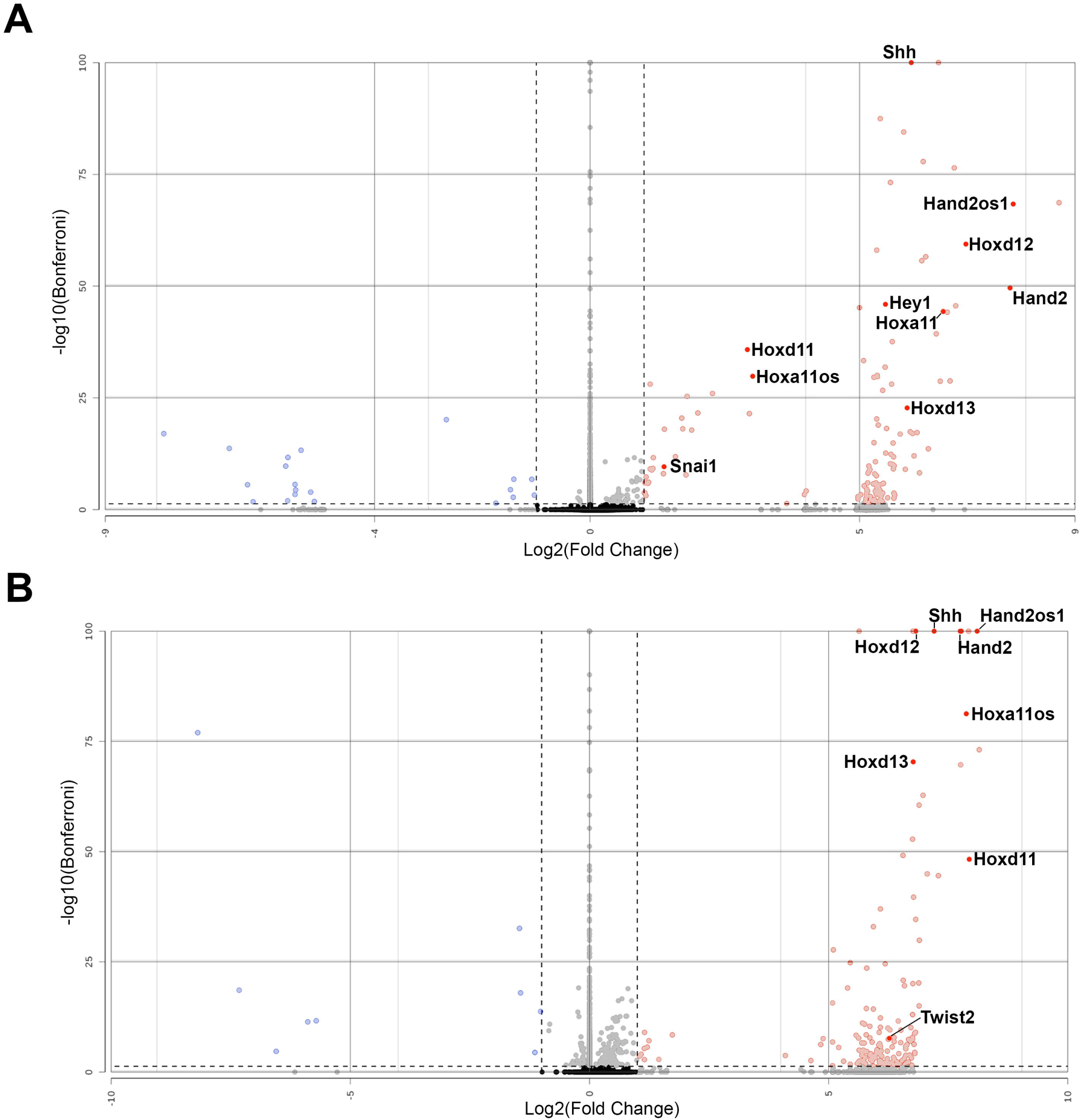
Differentially expressed genes in *Shh*-expressing cells. Volcano plots of differentially expressed genes (DEGs) detected in single-cell RNA-seq between Shh+ vs. Shh-mouse forelimb cells at e10.5 (A) and e11 (B) analyzed by Kruskal-Wallis. Left and right vertical dotted lines represent a ±2-fold change of expression in Shh+ cells compared to Shh-cells. The horizontal dotted line near the bottom of the graph represents the 0.05 Bonferroni-adjusted p-value cutoff.

### Of the Hox sites within the ZRS, Hoxd13 has its greatest affinity to the targeted Hoxd13 sites

Since Shh-expressing cells have several upregulated Hox TFs, we evaluated the potential Hox binding sites within the ZRS. We compared the relative affinities of all homeobox sites within the conserved ZRS to homeodomain TFs known to be expressed in the limb (Figure 6). Hoxd13 has a higher relative affinity for the two sites targeted in this study (Hoxd13 sites 1 and 2) than for any of the other potential Hox binding sites. Some of the other 5’ Hox TFs also favor these sites, suggesting these sites may provide a competitive mechanism to localize ZRS activity during limb development. The Hox5 paralogs (a/b/c), which play a role with Plzf in restricting Shh expression from the anterior limb (Xu et al., 2013), have high affinity for Lhx2 sites 2 and 1. It should also be noted that preliminary studies have recently associated these Lhx2 sites with long-range enhancer function (Bower et al., 2024).

**Figure 6:**
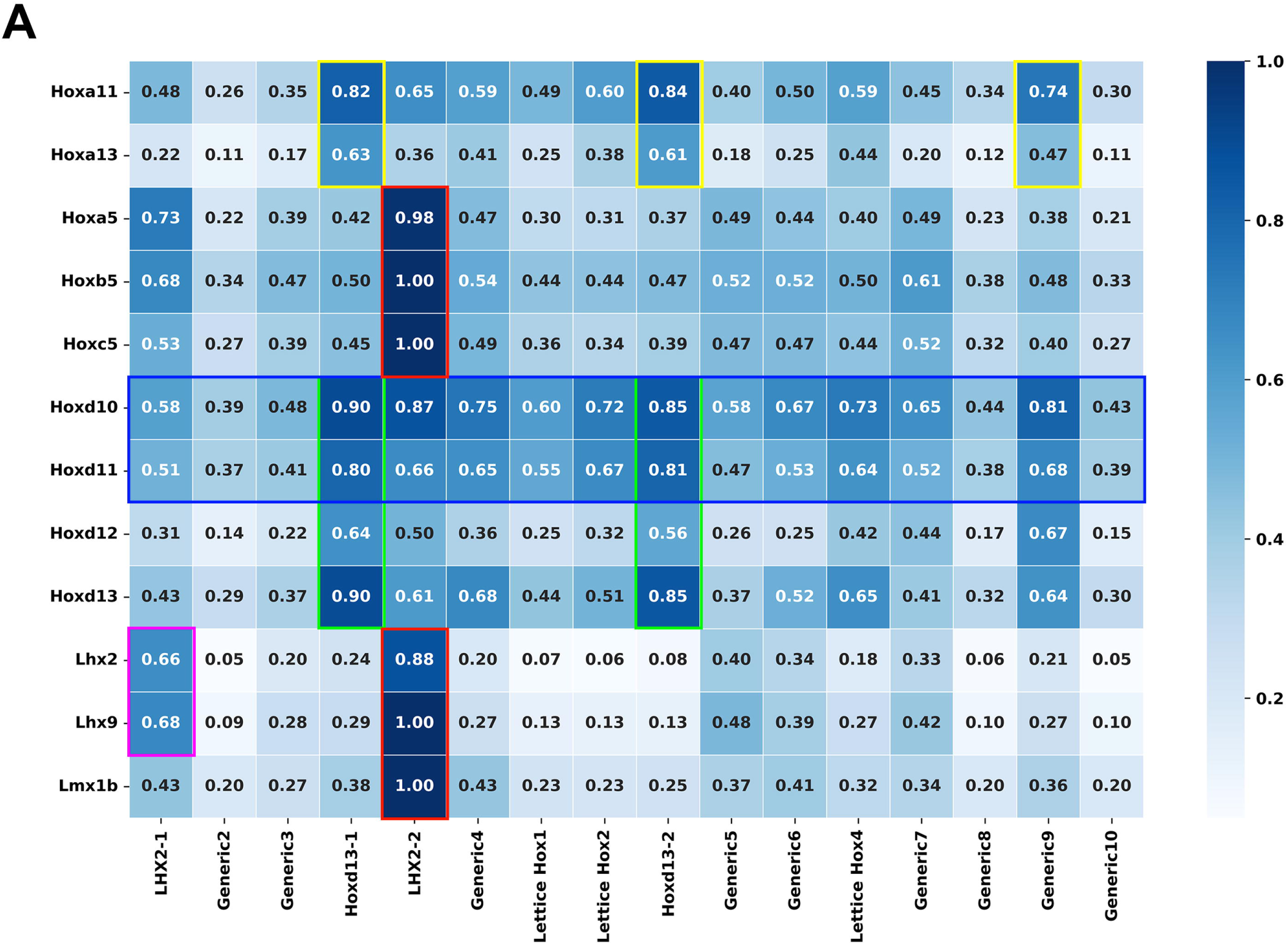
Hox Site Affinity. A heatmap relating the relative binding affinity of several homeodomain TFs (y-axis) to each possible homeobox sequence in the ZRS (x-axis). Affinity is expressed as a ratio on a scale of 0-1.

## 5 Discussion

The ZRS is an ∼800 bp highly conserved cis-regulatory module found within intron 5 of the *Lmbr1* gene (Lettice et al., 2003). Previous work has identified several important features of the ZRS including ETS and Hox binding sites that can be described as molecular rheostats controlling the relative level of ZRS activity (Lettice et al., 2012, 2017; Lim et al., 2024). The ZRS can be divided into three subdomains (5’, central, and 3’) based on the degree of sequence conservation (Figure 1). Our study also uncovered transcription factor binding sites that are critical for ZRS activity. Each of the three subdomains of the ZRS contain an E-box with the capacity to interact with basic helix-loop-helix transcription factors such as Hand2. We also evaluated two Hoxd13 binding sites, one in the 5’ subdomain and one in the central subdomain.

Disruption of the three E-boxes and two Hoxd13 binding sites (ZRSΔ5) nearly abates ZRS activity (Figures 2 & 4). By restoring each of the sites individually and in combination, we discovered that the 3’ E-box (E-box 3) and both Hoxd13 sites (1 and 2) can activate transcription, while the 5’ and central E-boxes (E-boxes 1 and 2) play inhibitory roles. Thus, we conclude that E-box 3 is the most likely site for Hand2 interaction.

The conserved 3’ subdomain (302bp) of the ZRS that was contained within our Fragment 3 (F3) is the only subdomain to retain activity, although it is not sufficient for full ZRS activity. Further, only constructs containing the 3’ subdomain have substantial activity (Figure 1) suggesting it contains binding sites that are necessary for initiation. Indeed, the 3’ subdomain contains three ETS binding sites, a Hox binding site, and an overlapping retinoic acid receptor (RAR)/NFκB/ETS4 site (Figure 5). In addition, we found a critical E-box in the 3’ subdomain (E-box 3) that promotes robust ZRS activity. The importance of this E-box was demonstrated when it was reintroduced into our ZRSΔ5 (ZRSΔ5+E3, Figure 3B) construct and recovered ZRS activity. However, E-box 3 is not essential as its absence does not prevent activity in the full-length ZRS when the functional Hoxd13 binding sites are present (ZRSΔ5+Hx1Hx2, Figure 3A). There is also an ETV binding site within the 3’ subdomain allowing ETV4/5 to recruit histone deacetylases to restrict chromatin access and subsequent activation (Lettice et al., 2012). Nevertheless, the individual and combined 5’ and central fragments (F1, F2, and F1F2) may have little or no activity because they lack E-box 3 and other initiating sites within the 3’ subdomain.

Hand2 has long been recognized as a critical upstream transcription factor for Shh. Our data suggest the 3’ E-box (E-box 3) is key in Shh activation and not the consensus Hand2 binding site found in the central subdomain (E-box 2). This is supported by evidence that deleting the canonical Hand2 binding site does not affect Shh expression, shown by Lettice *et al*. (2017). Osterwalder and colleagues demonstrated interaction between Hand2 and amplicons containing E-box 2 in e10.5 mice using ChIP-qPCR, though the 5’ E-box (E-box 1) and E-box 3 were not tested (Osterwalder et al., 2014). While these data show Hand2 can bind E-box 2, our data suggest that E-boxes 1 and 2 are repressive, not activating as was previously thought.

Basic helix-loop-helix transcription factors, such as Hand2, require dimerization. Koyano-Nakagawa *et al*. showed that Hand2 binds E-box 1 when heterodimerized with E47, but not as a homodimer (Koyano-Nakagawa et al., 2022). Further, Dai and Cserjesi showed that Hand2 can form homodimers, but that only Hand2-E12 heterodimers were transcriptionally active in yeast- and mammalian-two-hybrid systems (Dai and Cserjesi, 2002). Taken together this suggests that Hand2 is capable of binding E-boxes 1 and 2, but at these sites it may not play an activating role (Welscher et al., 2002).

Moreover, Hand2’s role could vary based on its dimerization partner (Firulli et al., 2007). Our single-cell analysis demonstrates upregulation of other E-box-binding factors such as Hey1, Snail, and Twist2 in Shh expressing cells suggesting a role in Shh regulation. It is also becoming increasingly clear that Hand2 is flexible enough to utilize E-boxes that differ from its consensus motif (Fernandez-Perez et al., 2019), which may depend upon its dimerization partner. Hand2’s ZRS-related dimerization partners remain to be determined.

The 5’ and central subdomains can contribute to ZRS activity but have little to no activity on their own. The 5’ subdomain has a Hoxd13 binding site, an E-box (E-box 1) and two ETS binding sites. These ETS binding sites (ETS0 & ETS1) are critical for overall ZRS activity and are missing in snakes (Kvon et al., 2016; Leal and Cohn, 2016). Interestingly, there is initial Shh expression in pythons that appears to coincide with the induction of Shh by ETV2 (Leal and Cohn, 2016; Koyano-Nakagawa et al., 2022) suggesting that ETS binding sites other than those of the 5’ subdomain can initiate expression but require these ETS sites to amplify or maintain Shh expression. Lettice and co-workers found several ETS binding sites within the 3’ subdomain; ETS3, which is less than 20 bp from E-box3, was important for full activity (Lettice et al., 2012). We found that E-box 1, which is near ETS1, inhibited ZRS activity. However, when a portion of the 5’ subdomain containing the Hoxd13 binding site, ETS0, and ETS1, but lacking E-box 1, was coupled to the 3’ subdomain (F1F3), it was sufficient to recover full ZRS activity (Figure 1) suggesting that a role for the 5’ region is to amplify the activity of the 3’ subdomain.

The central ZRS subdomain includes a recognized 5 bp inhibitory sequence identified from the human condition Werner mesomelic syndrome (WMS) and is tightly linked to the Hoxd13 site 2. This site has also been implicated by Xu and colleagues as an inhibitory site used by Hox5 paralogs to restrict the anterior activity of the ZRS through interaction with Plzf, whose binding site overlaps the WMS region (Xu et al., 2013). In our Hox binding site affinity analysis, we found that the Hox5 paralogs have a very high relative affinity (0.98-1.00) for a Hox site previously identified as an Lhx2 binding site, though they could also bind to Hoxd13 site 2 (Figure 6). In our studies we found that the central E-box (E-box 2) had an inhibitory effect on activity when present (Figure 4).

In addition, single nucleotide variations (SNVs) in the central ZRS often lead to anterior ectopic Shh expression; these have been linked to preaxial polydactyly, syndactyly, triphalangeal thumb, and WMS. Remarkably, the majority of clinically significant SNVs result in increased Shh expression either by loss of a repressor or gain of an activator (Bass et al., 2015) (Supplementary Table 1 and Supplementary Figure S1). Lim and co-workers demonstrated that subtle increases in ETS binding affinity could extend ZRS activity into the anterior margin (Lim et al., 2024) causing ectopic Shh expression and explaining some SNVs associated with preaxial polydactyly. Repressors such as Etv4 and Etv5 have been reported to inhibit the ZRS anteriorly and localize its expression to the ZPA (Lettice et al., 2017). There is an ETV binding site within the central region and within the 3’ subdomain. These ETVs are thought to recruit histone deacetylases restricting chromatin accessibility and ZRS activity. Taken together, these data suggest the central ZRS subdomain is dispensable for activity, may be important for localization, and tends to foster ectopic activity when disrupted.

Our data, combined with previous reports, support a working model of ZRS activity with three modules. First, the 3’ activation subdomain contains an E-box (the likely site of Hand2 binding) and ETS binding sites, of which at least one is likely required for initiation, and a binding site (identified as ETVB) that can toggle ZRS activity on or off depending on whether it is occupied by GABPα or ETV4/5, respectively. Second, the 5’ amplification subdomain contains two ETS binding sites (ETS0 & ETS1) that enhance activity, a Hoxd13 site that enhances activity, and an inhibitory E-box (E-box 1). Finally, the central localization subdomain contains inhibitory sequences, the WMS sequence, E-box 2, Hox5 paralog-Plzf interacting domain, and multiple enhancing Hox sites including a Hoxd13 site. These modules of the ZRS are represented in the diagram in Figure 7 and work collectively to initiate, maintain, and localize Shh expression to the posterior sub-AER mesoderm during limb outgrowth

**Figure 7.**
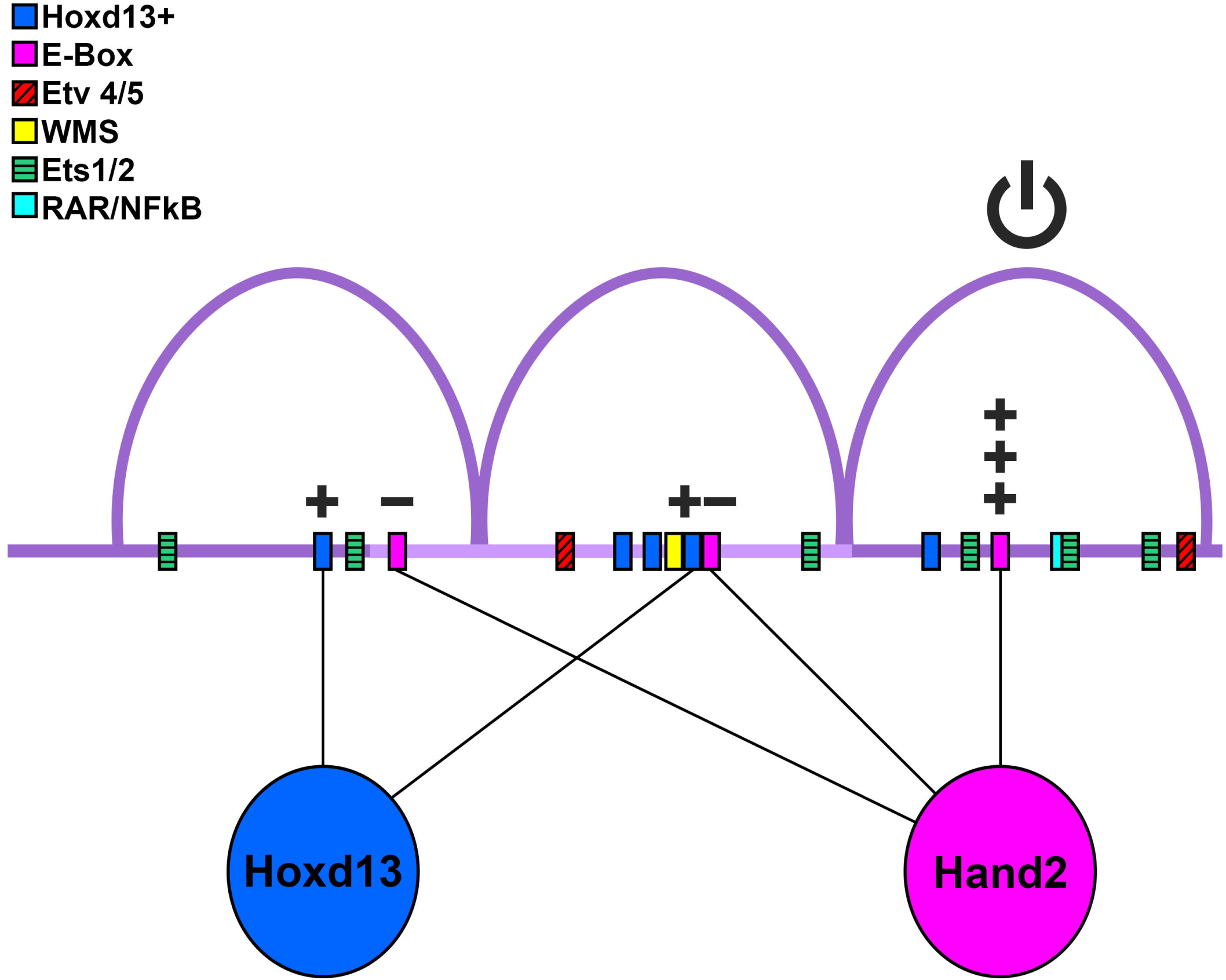
Model of binding site utilization. Diagram of ZRS with possible transcription factor-binding site interactions. Hoxd13 and its putative binding sites are shown in blue, Hand2 and E-boxes are depicted in pink, although other transcription factors may bind the E-boxes. Plus signs indicate an increase of ZRS activity, minus signs represent reduction of activity. The power (on/off) symbol represents that the 3’ conserved region is necessary for ZRS activity.

## 7 Statements

### Author Contributions

KFB: Writing – original draft, review, & editing, Conceptualization, Data curation, Formal analysis, Investigation, Methodology, Visualization.

SM: Writing – review and & editing, Conceptualization, Data curation, Formal analysis, Investigation, Methodology.

AKU: Writing – review and & editing, Investigation.

SRR: Writing – review and & editing, Investigation.

MMM: Writing – review and & editing, Investigation.

JA: Writing – review and & editing, Investigation.

AMC: Writing – review and & editing, Formal analysis, Investigation.

CUP: Writing – review and & editing, Conceptualization, Investigation, Methodology,

KCO: Writing – review and & editing, Conceptualization, Funding Acquisition, Resources, Visualization.

## Acknowledgements

The authors would like to thank Jessica Treto, Christopher G. Wilson, and the members of the Oberg Lab.

## Data Availability

Datasets are available on request:

The raw data supporting the conclusions of this article will be made available by the authors, without undue reservation. The original code used in this study is available at https://github.com/KateBall/Quantitative_Image_Analysis under the GNU Public License (GPL, ver. 3).

