## Supplementary Figures and Legends for "The 3’ Region of the ZPA Regulatory Sequence (ZRS) is required for activity and contains a critical E-box"

### *Supplementary Material*

#### 1 Supplementary Data

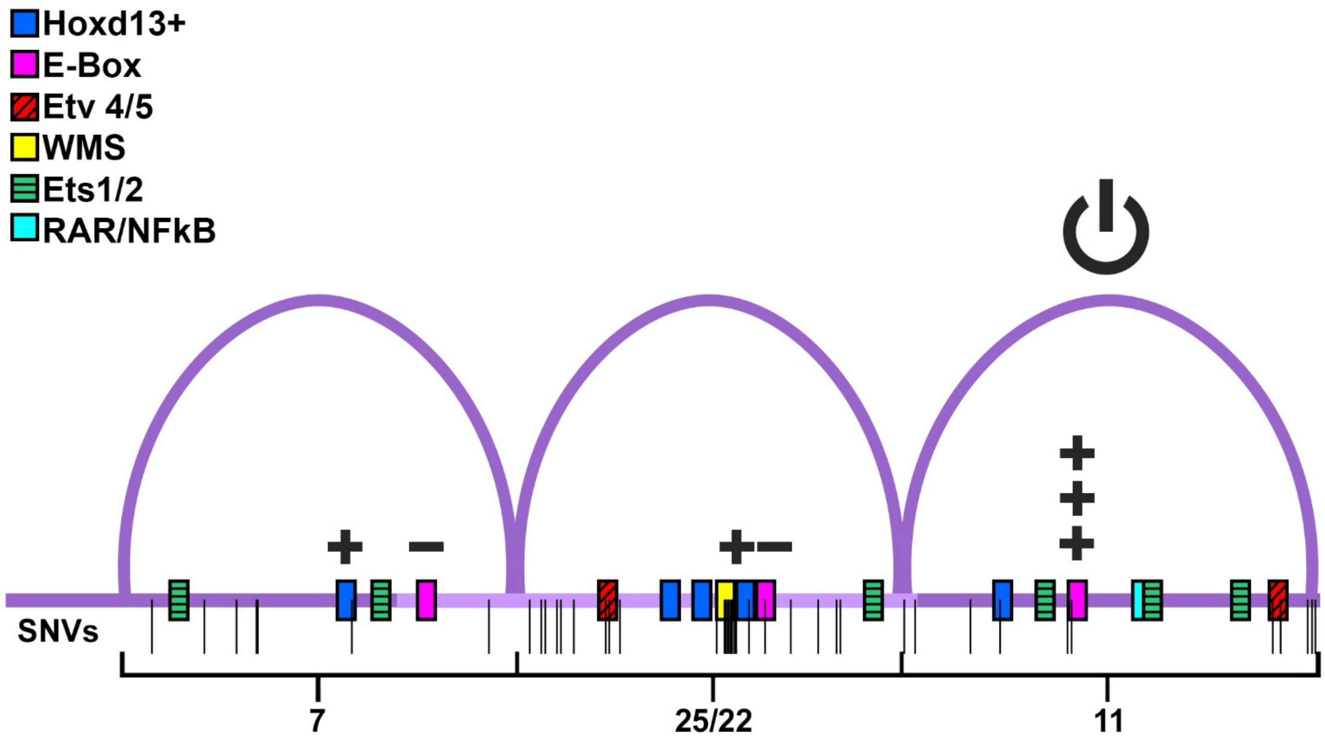


**Supplementary Figure 1: Distribution of Reported Single Nucleotide Variations (SNVs) within the ZRS Subdomains.** This is a graphic representation of SNVs within the ZRS that are reported to demonstrate preaxial polydactyly or ectopic activity in transgenic mice as listed in Supplementary Table 1. The SNVs are aligned on a schematic of the ZRS subdomains with predicted transcription factor binding sites. The central subdomain contains 25 disruptive SNVs that target 22 different sites, compared with 7 in the 5' subdomain and 11 in the 3' subdomain. The distribution suggests an important role for the central subdomain in localizing ZRS activity with the Werner-mesomelic syndrome (WMS) site being a hot spot for pathogenic SNVs (10 SNVs over 7 sites from n401-417) associated with triphalangeal thumbs and preaxial polydactyly. All 9 of the SNVs from this region that have been tested, demonstrated anterior ectopic ZRS activity in transgenic mice.

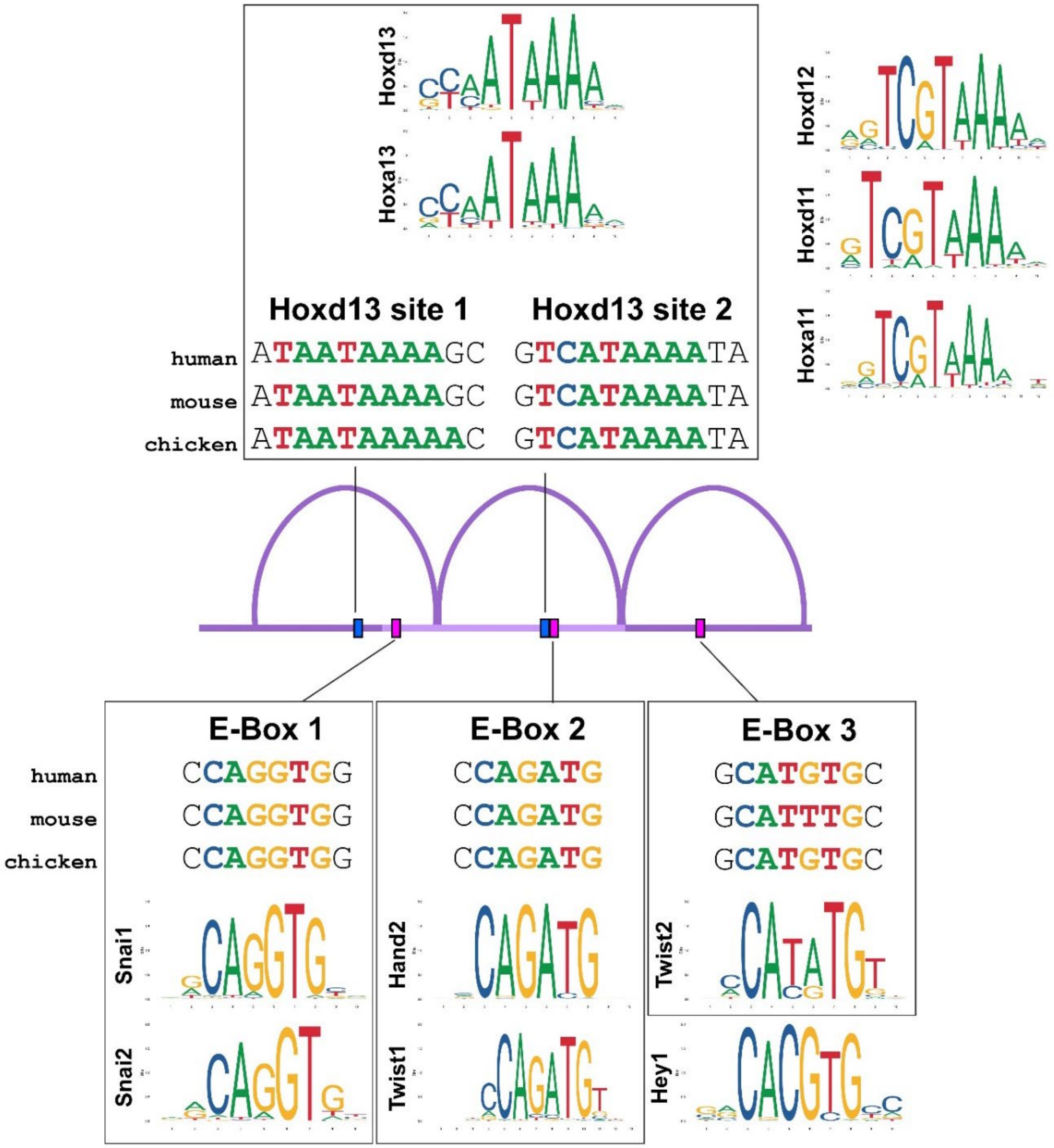


**Supplementary Figure 2: Transcription factor consensus binding motifs suggest possible regulators for each ZRS conserved binding site.** A ZRS diagram showing the Hox and E-box sequences relevant to this study. Top: Hoxd13's degenerate motif, [C/T][A/C]ATAAA, is found twice in the ZRS at sites we have called Hoxd13 site 1 and 2. It should be noted that Hoxa13's core binding motif is nearly identical to Hoxd13's motif, though Hoxa13 is expressed later in limb development. Other homeodomain factors that could act on the ZRS (Hoxa11, Hoxd11, Hoxd12) are less likely to use these sites due to their consensus motifs. Bottom: Each of the ZRS E-boxes bears a resemblance to the binding motifs of TFs known to be co-expressed with Shh in the limb. In addition to Twist 2's motif, E-box 3 is identical to sequence bound by Hand2/E12 (Dai and Cserjesi, 2002). For ease of comparison, the reverse complements of Hoxd13 site 2 and E-Box 3 are shown. Sequence Logos downloaded from JASPAR (https://jaspar.elixir.no/) (Rauluseviciute et al., 2023).

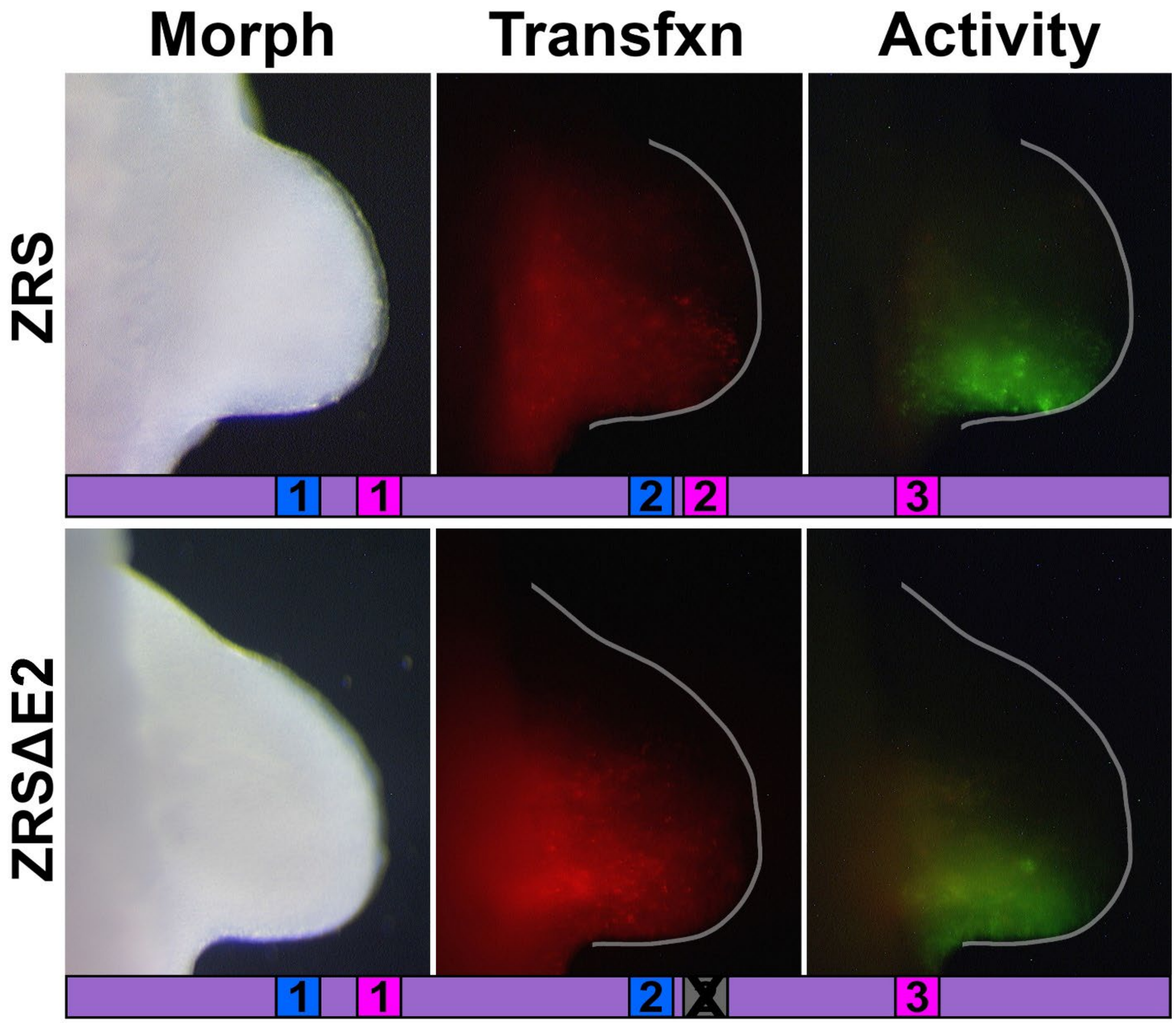


**Supplementary Figure 3: ZRS activity persists despite disruption of the central E-box 2 (E2).** Hand2 has been shown to bind to the E-box of the central subdomain, E-box 2 (E2), but activity of the ZRS persists even when the E2 binding site is disrupted. This finding is consistent with other reports that indicate that the central domain is not necessary for ZRS activity (Lettice et al., 2017).

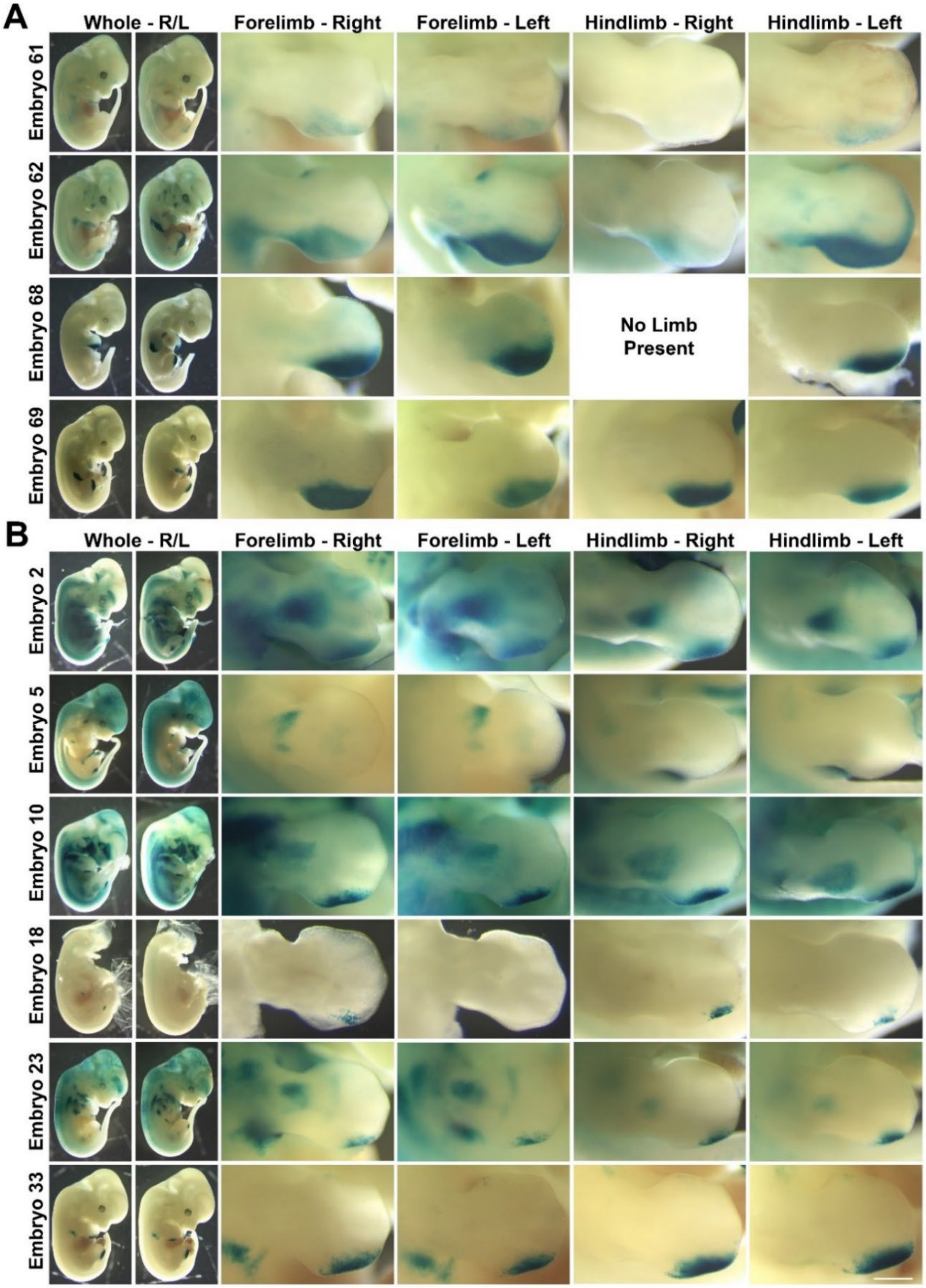


**Supplementary Figure 4: Transgenic Mouse Embryos.** A) Mouse embryos (e12.5) that were successfully transfected with wild-type hZRS in the HSP68-LacZ plasmid. B) Mouse embryos (e12.5) that were successfully transfected with HSP68-LacZ plasmid harboring hZRSΔ5. Scale bar represents 1mm.

**A**

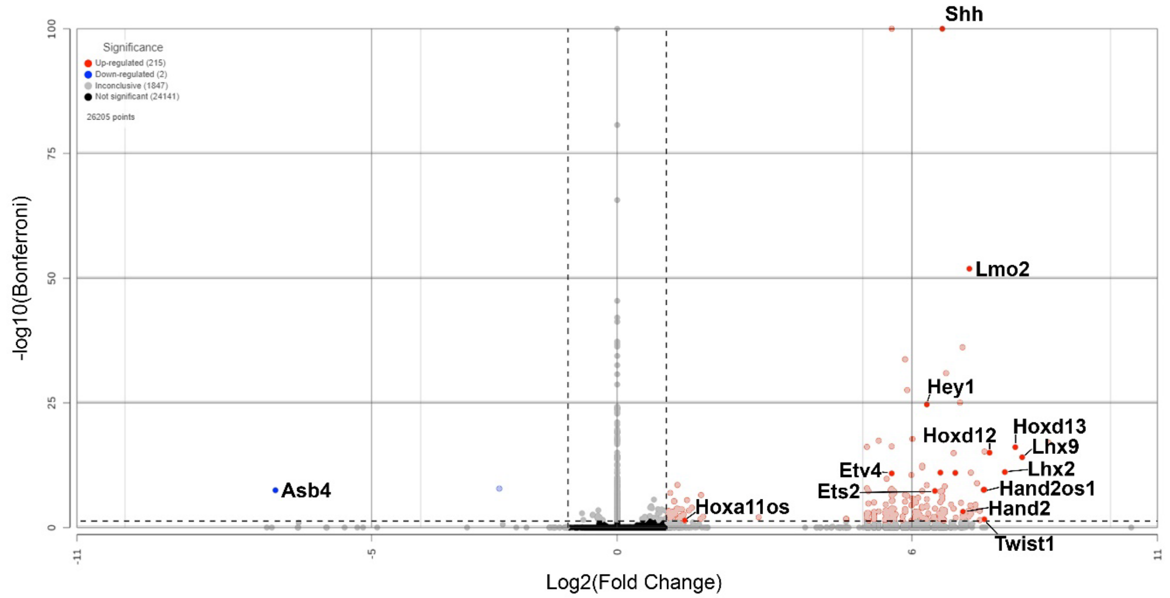


**Supplementary Figure 5: Differentially expressed genes in Shh-expressing cells at e12.** A) Volcano plot of differentially expressed genes (DEGs) detected in single-cell RNA-seq between Shh+ vs. Shh- mouse forelimb cells at e12 analyzed by Kruskal-Wallis. Left and right vertical dotted lines represent a ±2-fold change of expression in Shh+ cells compared to Shh- cells. The horizontal dotted line near the bottom of the graph represents the 0.05 Bonferroni-adjusted p-value cutoff.

### 2 Supplementary Tables

#### 2.1 Supplementary Table 1

**ZRS Single Nucleotide Variations (SNVs).** A table of all ZRS SNVs documented to date, including subdomain location, references, and phenotypes. CD: clinodactyly, ED: ectrodactyly, HF: hypoplastic fibula, HH: hypoplastic hallux, HP: hypoplastic pollex, HR: hypoplastic radius, HT: hypoplastic tibia, HU: hypoplastic ulna, PAP: postaxial polydactyly, PPD: preaxial polydactyly, SD: syndactyly, TH: thenar hypoplasia, TPT: triphalangeal thumb, WMS: Werner-mesomelic syndrome.

#### 2.2 Supplementary Table 2

**Primers used in this study.** All primers are listed and were made by Integrated DNA Technologies Inc., (Coralville, Iowa). Note that the cZRS construct was generated by first pulling-down a larger fragment that included the pre-ZRS with the listed primers, then a truncated version of was generated using the Erase-a-Base system.

#### 2.3 Supplementary Table 3

**Key Differentially Expressed Genes from Single-cell Analysis.** Mouse limb scRNA-seq data of Shh-related factors and potential regulators/cofactors that could act through the binding sites within this study. The log2 fold change in expression of Shh-expressing cells (Shh+) vs. Shh-non-expressing cells (Shh-), along with the Bonferroni-corrected p-value, for each gene is given for stages e10.5, e11, and e12. A threshold of ±2 fold-change (±1 log2 fold-change) was used to determine upregulated (red) and downregulated (blue) genes.

### 3 Chicken Embryo Limb Inclusion Criteria

#### 3.1 Description

Embryo limbs were ranked on a 0-5 scale for morphology, transfection based on RFP, and autofluorescence. Limbs needed to score a 3 or above in each category to be included in our analysis.

Morphology: TREP can cause significant damage to limb tissue making it difficult to assess transfection quality or enhancer activity pattern.

Transfection: Transfection coverage and quality can vary due to many factors involved in TREP such as subtle differences in injection location, DNA solution leaking out of the coelom before the current is applied, and differences in electrical conductivity during electroporation.

Autofluorescence: light in a fluorescent image that 1) can be outside the transfected region 2) can appear yellow-green in a GFP image and can result from an embryo that was dead prior to harvest, or an embryo that was not imaged quickly enough after harvest.

### 3.2 Limb Grading Rubric

| Score | 0 | 1 | 2 | 3 | 4 | 5 |
|---|---|---|---|---|---|---|
| **Morphology** | Limb is absent or indistinguishable from surrounding tissue | Limb is present but severely malformed or damaged from TREP or harvest, anatomical axes may be unclear. | Limb may be moderately malformed, anatomical axes are clear, limb is less than 50% the size of the contralateral limb. | Limb may be minimally malformed, anatomical axes are clear, limb is at least 50% the size of the contralateral limb. | Limb is fully formed, anatomical axes are clear, size is at least 75% of the contralateral limb, may have superficial defects. | Limb is fully formed, anatomical axes are clear, size is approximately equal to the contralateral limb, no obvious defects. |
| **Transfection efficiency** | No RFP is visible above background. | Only faint RFP is visible. | RFP is visible but may be faint or may not cover the target transfection region. | RFP is clearly visible and covers most of the target region. | RFP is bright and covers the entire target region | RFP is very bright and covers the entire target region |
| **Auto-fluorescence** | An extreme amount of autofluorescence is visible | An excessive amount of autofluorescence is visible | A moderate amount of autofluorescence is visible | A minimal amount of autofluorescence is visible | A very minimal amount of autofluorescence is visible | No autofluorescence is visible |

** Autofluorescence as just a commentary
