## Supplementary Tables and Legends for "The 3’ Region of the ZPA Regulatory Sequence (ZRS) is required for activity and contains a critical E-box"

Supplementary Table 1

**All ZRS SNVs documented to date**: including subdomain location, references, and phenotypes. CD: clinodactyly, ED: ectrodactyly, HF: hypoplastic fibula, HH: hypoplastic hallux, HP: hypoplastic pollex, HR: hypoplastic radius, HT: hypoplastic tibia, HU: hypoplastic ulna, PAP: postaxial polydactyly, PPD: preaxial polydactyly, SD: syndactyly, TH: thenar hypoplasia, TPT: triphalangeal thumb, WMS: Werner-mesomelic syndrome.

| Position | Change | Name | Organism | Subdomain | References | Phenotype | Kvon 2020 activity | Binding Sites Lost/Altered | Binding Sites Gained | Proposed Mechanism |
|---|---|---|---|---|---|---|---|---|---|---|
| 41 | A>C | NA | Human | 5' (less conserved) | Kvon 2020 | TPT, PPD, ED, HT | No impact | | | |
| 71 | T>C | US1011001 | Human | 5' | Kvon 2020 | PPD | No impact | Lhx2 site 1 | | |
| 93 | G>T | NA | Human | 5' | Pu et al. 2024 (preprint) | PPD | NA | | | |
| 105 | C>G | Dutch 1 | Human | 5' | Heutink 1994, Lettice 2003, Zhao 2016, Fuxman 2015, Amano 2017, Baas 2017, Cai 2019, Zhang 2019 | TPT, PPD, SD, PAP | Ant. Gain | | TFAP 2B/2A/2E, NF1 | Potential AP2 bs |
| 106 | G>A | NA | Human | 5' | Pu et al. 2024 (preprint) | PPD | NA | | | |
| 165 | A>G | Dutch 2 | Human | 5' | Lodder 2009, Potuijt 2020, Lim 2024 | TPT (mild, maybe subclinical) | No impact | GATA-1 | C/EBP Alpha | |
| 252 | G>C | UK Cat 1 | Cat | 5' | Lettice 2008 | PPD | No impact | | | |
| 278 | G>A | NA | Human | Central | Pu et al. 2024 (preprint) | PPD | NA | | | |
| 285 | G>T | Silkie 1 (Gal) | Chicken | Central | Dorshorst 2010, Amano 2017 | PPD | No impact | Irx 3/5 | | Repressor lost |
| 287 | C>A | Balochi or Pakastani 1 | Human | Central | VanderMeer 2012, Dai 2013 | TPT, PPD, PAP, SD | Ant. Gain | Irx 3/5 | | Irx3/5 bs |
| 295 | T>C | UK low-penetrant; Dutch 3 | Human | Central | Furniss 2008, Lodder 2009, Potuijt 2020, (Lettice 2003 reported this, but was found in unaffected individuals) | TPT, PPD, CD, TH | No impact | | | |
| 297 | G>A | French 1 | Human | Central | Albuisson 2011 | TPT, PPD | No impact | SP-1 | | |
| 305 | A>T | Belgian 1 | Human | Central | Lettice 2003 | TPT, PPD | No impact | SP-1 | SP-1 | ETS1 |
| 328 | C>G | Indian 2 | Human | Central | Kvon 2020, Lim 2024 | TPT, PPD, TH | No impact | ETV4/5 | | |
| 329 | T>C | Belgian 2 | Human | Central | Lettice 2003, Lettice 2008, Dai 2013, Kvon 2020 | TPT, PPD | No impact (except 1/5) | ETV4/5 | Pea3 | ETS1 |
| 334 | T>G | French 2 | Human | Central | Albuisson 2011, Lim 2024 | PPD | Ant. Gain | C/EBP beta | | |
| 396 | C>T | Turkish 1 | Human | Central | Semerci 2009, Kvon 2020, Francisco Gonzalez Alvarez 2022 | TPT, PPD, SD | Ant. Gain | ER, T3R-Alph | GR | |
| 401 | A>G | French 4 | Human | Central | Kvon 2020 | TPT, PPD, TH | Ant. Gain | Plzf | | |
| 401 | A>C | French 5 | Human | Central | Kvon 2020 | TPT, PPD, TH, HT | Ant. Gain | Plzf | | |
| 402 | C>T | Mexican | Human | Central | VanderMeer 2014 | WMS, TPT, PPD, HR, HT | Ant. Gain | ER, Plzf | | |
| 403 | T>C | NA | Human | Central | Zepeda-Olmos 2024 | TPT, PPD, TH | NA | Plzf | | |
| 404 | G>T | Indian 1 | Human | Central | Lettice 2008, Girisha 2014, Al-Qattan 2018 | WMS, TPT, PPD, HR, HU, HT | Ant. Gain | Plzf | C/EBP Alpha | |
| 404 | G>C | Brazilian; French 6 | Human | Central | Wieczorek 2010, Kvon 2020 | WMS, TPT, PPD, HH, HT, HF | Ant. Gain | Plzf | | |
| 404 | G>A | Cuban; Turkish 2; Korean | Human | Central | Zguricas et al 1999, Lettice 2003, Wieczorek 2010, Norbnop et al 2014, Cho 2013, Lettice 2017 | WMS, TPT, PPD, HT, HF | Ant. Gain | Plzf | | |
| 406 | A>G | M100081 (Mus); Thai | Mouse; Hum | Central | Masuya 2007, Norbnop 2014, Amano 2017 | WMS, TPT, HT | Ant. Gain | Plzf/Hoxd13 site 2 | C/EBP Beta | Repressor lost |
| 407 | T>A | DZ Mouse; 5460001 | Mouse; Hum | Central | Zhao 2009, Kvon 2020 | TPT, PPD, HT | Ant. Gain | | | |
| 417 | A>G | Mosaic or French 3 | Human | Central | Vanlerbergh 2015, Al-Qattan 2018 | TPT, PPD, HT | Ant. Gain | c-Jun, p40x, E-Box | E-Box becomes D-Box | Hand2 bs |
| 428 | T>A | Chinese 1 | Human | Central | Wu 2016 | TPT, PPD | No impact | c-Jun, GCN4, p40x | SP-1 | |
| 446 | T>A | Chinese 2 | Human | Central | Xu 2020 | PPD | NA | Oct-1, Oct-6 | | HnRNP K |
| 463 | T>G | Pakastani 2 | Human | Central | Farooq 2010 | TPT, PPD | Ant. Gain | | | |
| 475 | A>G | Hemmingway Cat | Cat | Central | Lettice 2008 | PPD | No impact | | | |
| 477 | A>T | UK Cat 2 | Cat | Central | Lettice 2008 | TPT, PPD | No impact | | | |
| 507 | C>G | French 7 | Human | 3' | Kvon 2020 | TPT, PPD | No impact | | | |
| 515 | C>A | Silkie 2 (Gal) | Chicken | 3' | Maas et al 2011; Dunn et al 2011 | PPD | Ant. Gain | | | |
| 555 | G>A | Hx (Mus) | Mouse | 3' | Lettice 2008, Amano 2017, Lettice 2017 calls it 553, Mass 2005 | PPD | Ant. Gain | | | New MSX1 bs |
| 574 | G>A | French 8 | Human | 3' | Kvon 2020 | TPT, PPD | No impact | | | |
| 619 | C>T | Saudi | Human | 3' | Al-Qattan 2012, Al-Qattan 2018 | TPT, PPD, SD, LPAD, HP, HR | Ant. Gain | C/EBP Beta | C/EBP Alpha, USF | |
| 621 | C>G | US Family B | Human | 3' | Dobbs 2000, Gurnett 2007 | TPT, PPD | Ant. Gain | | | Cdx Binding sites |
| 739 | A>G | US Family AC | Human | 3' | Gurnett 2007, Lettice 2012 | TPT, PPD | Ant. Gain | | ETS1 | |
| 743 | T>G | Australian | Human | 3' | Lettice 2012 | TPT, PPD, TH | Ant. Gain | ETV4/5 | | |
| 763 | T>G | NA | Human | 3' (less conserved) | Kvon 2020 | NA | Ant. Gain | Sox5/6/9 bs | | |
| 767 | G>A | NA | Human | 3' (less conserved) | Kvon 2020 | NA | Ant. Gain | Sox5/6/9 bs | | |
| 769 | T>C | M101116 (Mus) | Mouse | 3' (less conserved) | Masuya 2007, Lettice 2008, Kozhemyakina 2014, Kvon 2020 | PPD | Ant. Gain | Sox5/6/9 bs | | GATA6 normally represses |

Supplementary Table 2

**Primers used in this study:** All primers are listed and were made by Integrated DNA Technologies Inc., (Coralville, Iowa). Note that the cZRS construct was generated by first pulling-down a larger fragment that included the pre-ZRS with the listed primers, then a truncated version of was generated using the Erase-a-Base system.

| Purpose/Mutation | Fwd Primer | Fwd Primer Seq 5'-3' | Rev Primer | Rev Primer Seq 5'-3' | Introduced RE cut site |
|---|---|---|---|---|---|
| cZRS (and Pre-ZRS) Genomic Primers | LS_SH_5p2 | CCAACCACTTGCATATTATGC | ZRS 3p2 | GACATCAAGGAATGACAAAGC | NA |
| cZRS Hoxd13 site 1 | ZRS Hoxd13 B1 SDM 5p1 | GCATGATAACTAGTACAAATAGTACAAAAATTTGAGG | NA | NA | SpeI |
| cZRS Hoxd13 site 2 | ZRS Hoxd13 B2 SDM 5p1 | CCCTGTACTGTATCTAGAGACCAGATGAC | NA | NA | XbaI |
| cZRS Hoxd13 site 2 and E-box 2 | ZRS Hand2 Hoxd13 B2 SDM 5p1 | CTGTATCTAGAGACTTAAGGACTTTTTTCCC | ZRS Hand2 Hoxd13 B2 SDM 3p1 | GGGAAAAAAGTCCTTAAGTCTCTAGATACAG | XbaI, AflII |
| cZRS E-Box 1 | ZRS Twist1 B1 SDM 5p1 | GGTAGACCTTAATTAAGCGAAGAGGCC | NA | NA | PacI |
| cZRS E-Box 2 | ZRS Hand2 SDM 5p2 | CTGTATTTTATGACTTAAGGACTTTTTTCCC | NA | NA | AflII |
| cZRS E-Box 3 | ZRS Twist1 B3 SDM 5p1 | GCTTAGTGTTAGTGGTTTAAACACATTCTTGG | NA | NA | PmeI |
| hZRS Genomic Primers | hZRS 5p1 | TTCTGCAGTATGTGGCTC | hZRS 3p1 | CTTGAAGGTGTTGGGAAAATC | NA |
| hZRS Hoxd13 site 1 | hZRS Hoxd13 B1 SDM 5p1 | GCCTGATACCCGGGGCAAAAAGTACAAAATTTTAGG | hZRS Hoxd13 B1 SDM 3p1 | NA | SmaI |
| hZRS Hoxd13 site 2 and E-box 2 | hZRS Hand2 Hoxd13 B2 SDM 5p2 | GTATCTAGAGACTTAAGGACTTTTTCCCCCCAGTGGC | hZRS Hand2 Hoxd13 B2 SDM 3p1 | GTCCTTAAGTCTCTAGATACAGTACAAGGTCAC | XbaI, AflII |
| hZRS E-Box 1 | hZRS Twist1 B1 SDM 5p1 | GACTGACCGATATCAAGCGAAGAGTTCTGTGC | NA | NA | EcoRV |
| hZRS E-Box 3 | hZRS Twist1 B3 SDM 5p1 | GCTTAGTGTTAGTGGTTTAAACGCATATTTGGC | NA | NA | PmeI |

Supplementary Table 3

**Key Differentially Expressed Genes from Single-cell Analysis**: Mouse limb scRNA-seq data of Shh-related factors and potential regulators/cofactors that could act through the binding sites within this study. The log2 fold change in expression of Shh-expressing cells (Shh+) vs. Shh-non-expressing cells (Shh-), along with the Bonferroni-corrected p-value, for each gene is given for stages e10.5, e11, and e12. A threshold of ±2 fold-change (±1 log2 fold-change) was used to determine upregulated (red) and downregulated (blue) genes.

| Gene/Transcript | Group | Log2 Fold change (e10.5 SHH+ vs e10.5 SHH-) | Bonferroni (e10.5 SHH+ vs e10.5 SHH-) | Log2 Fold change (e11 SHH+ vs e11 SHH-) | Bonferroni (e11 SHH+ vs e11 SHH-) | Log2 Fold change (e12 SHH+ vs e12 SHH-) | Bonferroni (e12 SHH+ vs e12 SHH-) |
|---|---|---|---|---|---|---|---|
| Shh | Shh | 5.96 | 0.00E+00 | 7.21 | 0.00E+00 | 6.62 | 0.00E+00 |
| Hoxa11os | Homeobox | 3.02 | 1.47E-30 | 7.88 | 5.37E-82 | 1.36 | 2.47E-02 |
| Hoxa11 | Homeobox | 6.55 | 4.74E-45 | 0.00 | 4.39E-16 | NS | NS |
| Hoxa13 | Homeobox | 0.00 | 8.60E-05 | 0.00 | 5.11E-35 | 0.00 | 7.54E-15 |
| Hoxd9 | Homeobox | NS | NS | NS | NS | NS | NS |
| Hoxd10 | Homeobox | 1.17 | 2.34E-12 | 6.25 | 3.84E-08 | NS | NS |
| Hoxd11 | Homeobox | 2.92 | 1.63E-36 | 7.94 | 5.31E-49 | NS | NS |
| Hoxd12 | Homeobox | 6.97 | 4.14E-60 | 6.82 | 2.02E-114 | 7.58 | 9.82E-16 |
| Hoxd13 | Homeobox | 5.88 | 1.77E-23 | 6.77 | 4.28E-71 | 8.11 | 7.63E-17 |
| Lhx2 | Homeobox | 2.00 | 2.38E-22 | 7.76 | 1.98E-70 | 7.89 | 7.55E-12 |
| Lhx9 | Homeobox | NS | NS | 6.71 | 1.03E-07 | 8.25 | 8.04E-15 |
| Hand2 | E-box binding related | 7.79 | 2.56E-50 | 7.75 | 2.11E-117 | 7.04 | 6.89E-04 |
| Hand2os1 | E-box binding related | 7.85 | 4.55E-69 | 8.11 | 3.55E-186 | 7.46 | 2.83E-08 |
| Hey1 | E-box binding related | 5.48 | 1.20E-46 | 0.00 | 2.49E-69 | 6.30 | 2.09E-25 |
| Twist1 | E-box binding related | NS | NS | 0.54 | 1.94E-03 | 7.47 | 2.19E-02 |
| Twist2 | E-box binding related | 0.93 | 1.87E-02 | 6.27 | 2.26E-08 | NS | NS |
| Snai1 | E-box binding related | 1.37 | 2.54E-10 | 0.00 | 2.95E-16 | NS | NS |
| Snai2 | E-box binding related | NS | NS | NS | NS | NS | NS |
| Hand1 | E-box binding related | NS | NS | NS | NS | NS | NS |
| Tfap2c | Limb/Development related | 6.46 | 0.00E+00 | 0.00 | 1.58E-184 | NS | NS |
| Spry1 | Limb/Development related | 6.63 | 7.26E-45 | 0.00 | 2.26E-38 | NS | NS |
| Tbx3 | Limb/Development related | 6.42 | 4.80E-40 | 0.00 | 2.09E-37 | NS | NS |
| Elk3 | Limb/Development related | 5.64 | 1.51E-10 | 0.00 | 5.52E-09 | NS | NS |
| Bmp4 | Limb/Development related | 8.70 | 2.26E-69 | 7.93 | 9.75E-198 | 7.20 | 9.88E-12 |
| Ptch1 | Limb/Development related | 5.75 | 1.27E-17 | 5.10 | 1.97E-28 | 5.60 | 3.47E-02 |
| Lmo2 | Limb/Development related | 6.76 | 3.51E-77 | 6.76 | 2.78E-201 | 7.17 | 1.28E-52 |
| Dusp6 | Limb/Development related | 1.88 | 1.60E-18 | 7.30 | 2.75E-45 | 6.62 | 2.22E-06 |
| Ets2 | Limb/Development related | 1.70 | 3.37E-21 | 6.78 | 2.03E-40 | 6.47 | 4.59E-08 |
| Lef1 | Limb/Development related | 1.38 | 1.01E-18 | 6.56 | 6.93E-50 | 6.67 | 5.29E-09 |
| Etv5 | Limb/Development related | 5.31 | 3.08E-04 | 0.00 | 3.45E-15 | 5.59 | 3.12E-07 |
| Fzd10 | Limb/Development related | 6.23 | 2.72E-57 | 0.00 | 2.55E-63 | 5.91 | 2.72E-28 |
| Etv4 | Limb/Development related | 1.11 | 8.68E-29 | 0.00 | 1.84E-28 | 5.59 | 1.41E-11 |
| Meis1 | Limb/Development related | 0.00 | 3.94E-21 | 0.00 | 4.65E-20 | NS | NS |
| Col1a2 | Limb/Development related | NS | NS | 0.00 | 2.84E-04 | NS | NS |
| Shox2 | Limb/Development related | NS | NS | NS | NS | NS | NS |
| Gli3 | Limb/Development related | -1.75 | 3.61E-02 | 0.00 | 3.24E-04 | NS | NS |
| Meis2 | Limb/Development related | -6.70 | 2.04E-14 | 0.00 | 8.52E-28 | NS | NS |
| Pbx1 | Limb/Development related | -1.42 | 1.56E-07 | -1.44 | 1.12E-18 | NS | NS |
| Gas1 | Limb/Development related | -7.91 | 1.01E-17 | -7.33 | 2.79E-19 | NS | NS |
| Asb4 | Limb/Development related | -2.67 | 7.56E-21 | -8.19 | 1.03E-77 | -6.96 | 3.32E-08 |

Upregulated

Downregulated

NS and/or |Log2FC|<1
